# The structural basis of the EPCR-APC complex induced biased PAR1 signaling

**DOI:** 10.1101/2023.02.07.527434

**Authors:** Alexei Iakhiaev

## Abstract

Activated Protein C (APC) is an effector enzyme of the natural anticoagulant pathway. In addition to its anticoagulant function, endothelial protein C receptor (EPCR)-bound APC induces biased protease-activated receptor type 1 (PAR1)-mediated signaling. Despite intensive investigation, the mechanism of biased signaling is not completely clear. To gain new insights into APC-induced PAR1-biased signaling we reviewed the published data and created three-dimensional models of the proteins and their complexes involved in the early stages of PAR1 signaling. A comparative study of models related to canonical and biased signaling demonstrated that interactions between APC, EPCR, PAR1, and Caveolin-1 (Cav1) can provide plausible explanations for the differences between the two types of PAR1 signaling. The model suggests that the interaction of the PAR1 peptide 22-ARTRARRPESK-32 with 162-helix of APC positions the PAR1 N-terminus for the preferential cleavage at R46. By contrast, the hirudin-like sequence of PAR1 is involved in the positioning of the N-terminus of PAR1 for cleavage at R41 by thrombin in canonical signaling. The model and molecular dynamics (MD) simulations of the tethered ligand (TL) interaction with APC suggest that the TL facilitates direct interaction of the EPCR transmembrane (TM) domain with the PAR1 TM helices 6 and 7 by transient binding to the light chain of APC and keeping EPCR-APC in close proximity to PAR1. The biased signaling paradigm considers the ligand-induced conformational changes in PAR1 as solely being responsible for the biased signaling. Our models suggest that Cav1, EPCR, and PAR1 interactions can provide a selective advantage to biased signaling over canonical signaling. First, the complex comprised of caveolin-1 oligomer-EPCR-APC-PAR1 positions EPCR-APC and PAR1 at a distance favorable for PAR1 activation. Second, the Cav1 presence favors selectivity for the PAR1 bound β-arrestin-2, not the PAR1-bound G protein alpha (Gα) subunit. The potential reason for β-arrestin-2 selectivity includes Gα binding to the Cav1 and its immobilization resulting in the inability of PAR1-bound Gα to periodically interact with the plasma membrane required for its function. MD simulations of the PAR1-EPCR-β-arrestin-2 complex demonstrated that one of the mechanisms of the APC-induced PAR1-biased signaling is the interaction of the EPCR TM domain with the PAR1-bound β-arrestin-2, leading to the stabilization of the PAR1-β- arrestin-2 complex and activation of β-arrestin-2. Thus, models suggest that Cav1 and EPCR-APC mediated interactions provide a selective advantage for the β-arrestin-2 dependent biased signaling, not the G proteins mediated canonical signaling by the PAR1 receptor.

**Author summary:** The APC-biased PAR1 signaling in endothelial cells results in the barrier protection response while thrombin-induced PAR1 canonical signaling results in a pro-inflammatory response with endothelial barrier dysfunction. It has been demonstrated that caveolar localization and occupancy of the EPCR are required for APC-biased signaling, however, the molecular mechanism remained incompletely clear. Computational modeling of the structure of the signaling complex and its molecular dynamics simulations allowed us to propose plausible mechanistic explanations for the requirement of caveolin 1 for biased signaling. The models that assume direct binding of transmembrane domains of EPCR and PAR1 in the signaling complex allowed us to gain new insights into APC-biased PAR1 signaling and better understand the requirement of EPCR occupancy for biased signaling.

## Introduction

Signaling by G protein coupled receptors (GPCR) can be divided into canonical and biased signaling. An example of canonical GPCR signaling is thrombin-induced signaling via PAR1 located on the platelets and endothelial cells (EC). Thrombin activation of platelets through PAR1 induces the release of α-granules and the formation of platelet aggregates. Thrombin cleavage of PAR1 in ECs leads to exocytosis of von Willebrand factor (VWF) and other mediators from the Weibel–Palade bodies and a decreased endothelial barrier function that promotes leukocyte extravasation (1). PAR1 cleavage through R41 and R46 sites can engage distinct subsets of signaling molecules to initiate two different signaling pathways, termed canonical and biased signaling, respectively. An example of biased PAR1 signaling through cleavage of R46 is activated protein C (APC)-induced signaling in ECs. In contrast to canonical PAR1 signaling through cleavage of R41 that results in barrier-disruptive and pro-inflammatory effects, biased signaling results in improved endothelial barrier function and gene expression patterns corresponding to anti-inflammatory effects (2, 3). To better understand the mechanism of PAR1-biased signaling, we investigated the structure and dynamics of proteins that initiate APC- induced PAR1 signaling. To gain new mechanistic insights into biased signaling, we compared proteins and their complexes responsible for the initiation of canonical and biased signaling.

Thrombin is generated as a result of the activation of the blood clotting cascade after vascular injury and is the main enzyme responsible for clot formation to stop bleeding. An important stage in this process is the activation of PAR1, a thrombin receptor that is mostly expressed in platelets and vascular ECs (4). Thrombin is an effector enzyme of the canonical pathway which includes PAR1, G proteins, and other molecules downstream in this signaling pathway (1, 5). The APC represents an effector enzyme of the biased signaling pathway and is generated from zymogen protein C (PC) by the thrombin-thrombomodulin complex. The PC pathway is a clinically important natural anticoagulant pathway that influences coagulation, fibrinolysis, and inflammation (6). This pathway includes PC, cell surface thrombin receptor thrombomodulin, and endothelial PC receptor (EPCR). APC bound to EPCR activates the PAR1 and initiates biased signaling. In addition to APC, EPCR, and PAR1, other proteins participating in the initiation of the biased signaling pathway include Cav1, β-arrestin-2, and proteins downstream in the signaling pathway (2,3,7,8). In addition to being a receptor for thrombin, APC, and other proteases, PAR1 can serve as the sensor of shear stress forces in laminar flow or stretch (9).

Caveolae are considered sensors for the tensional forces in the cell membrane (10) and β- arrestins are important for mechanotransduction (11–13). APC-induced PAR1-biased signaling has several different cellular outcomes. The first outcome includes cytoprotective and anti- inflammatory effects via inhibition of gene expression induced by the NF-kB signaling pathway. The second outcome includes a block of the apoptotic pathways. The EPCR-APC-mediated PAR1 activation blocks both the mitochondria-mediated, and death receptor-mediated apoptotic pathways through the inhibition of caspase-8 (1). The third outcome includes the induction of the release of extracellular vesicles. Exposure of human umbilical vein endothelial cells or monocytes to APC results in the release of EPCR-containing microparticles (14, 15). The phenomenon requires the APC active site, EPCR, and PAR1 expressed on ECs (be consistent if you want to abbreviate the term). The FVIIa treatment induces the release of extracellular vesicles from the endothelium via FVIIa-EPCR-PAR1 mediated biased signaling (7). The biased signaling by the EPCR-APC-PAR1 is still poorly understood. It was demonstrated that the PAR1 bound β-arrestin-2 is responsible for the APC cytoprotective signaling (2,3,16), while signaling through G proteins leads to a proinflammatory response.

The puzzling observation demonstrated that occupancy of the EPCR by its ligands leads to the signaling through the PAR1- β-arrestin-2 pathway, independent of the cleavage site (2).

Studies with the APC-FVIIa chimera demonstrated that APC EGF-like 1 domain is important for biased signaling by the EPCR-APC-PAR1 system. The localization of PAR1 in caveolar or lipid raft microdomains is critical for APC-promoted biased signaling. It was shown that β-arrestin- 2 could be detected in Cav1 containing fractions in control cells (16, 17). The β-arrestins can be co-immunoprecipitated with PAR1 from unstimulated cells (3). The Cav1 can be immunoprecipitated with anti-PAR1 and anti-EPCR antibodies before stimulation of the cells with APC while Cav1 does not co-immunoprecipitate with the EPCR and PAR1 after treatment of the cells with APC (18). The three-dimensional structure of the APC-PAR1 signaling complex is not known. To generate new insights into the mechanism of APC-induced biased PAR1 signaling, we modeled the proteins and their complexes known to be involved in the biased signaling at early stages. The proteins include APC, EPCR, Cav1 oligomer, PAR1, and β- arrestin-2. We investigated the quaternary signaling initiation complex composed of APC- EPCR-PAR1-Cav1 that is responsible for TL formation and PAR1 activation. Then we investigated the PAR1-EPCR-β-arrestin-2 complex responsible for β-arrestin-2 activation and biased signaling. This study is designed as a comparative study of canonical and APC-biased PAR1 signaling that is based on the review of published data and creating models that are consistent with the experimental observations.

## Results and Discussion

### Proteins and protein complexes that are involved in the early stages of APC-mediated biased PAR1 signaling

The main participants of the early stages of APC-induced signaling include EPCR, PAR1, Cav1 oligomer, and β-arrestin-2. We divided this literature review complemented by computational studies into four sections. The focus of the first section is on building full-length structural models of the participating proteins and their complexes. This section investigates details of EPCR-APC complex interaction with the PAR1 N-terminus. The next section focuses on the interactions of Cav1 with PAR1 and EPCR. The third section focuses on PAR1 activation, and the fourth section on the activation of β-arrestin-2 and the EPCR-PAR1-β- arrestin-2 complex.

## 1. Structure, function, and dynamics of APC, EPCR, APC-EPCR complex

### 1.1 Interaction of APC with the PAR1 N-terminus

The structure of APC contains a light chain composed of a Gla-domain, two EGF-like domains, and a heavy chain containing the catalytic domain. The two-chain molecule is held together by a disulfide bond. The experimental three-dimensional structure of human Gla-domainless APC (PDB ID: 1aut.pdb) has been published (43). The published structure of APC does not contain the Gla-domain, a nine amino acids long C-terminal fragment of light chain, and ten C-terminal amino acids of the protease domain. We assembled a structure of full-length APC by adding atomic coordinates for missing parts to the experimental structure contained in 1aut.pdb. Atomic coordinates for missing parts were taken from the AlphaFold2 database (model ID: AF-P04070-F1-model_v2.pdb) and from the EPCR-APC Gla-domain complex (PDB ID: 1lqv.pdb). MD simulation of the APC was used to select the conformation of flexible parts of full-length APC. The ANM analysis demonstrated that the APC light chain contains several flexible regions. In addition to the light and heavy chain C-terminal peptides not resolved in crystal structure due to their flexibility, the regions between Gla- and the EGF1 domains as well as between EGF1 and EGF2 domains were also flexible and allow the molecule to adjust the active site’s position. To describe these motions quantitatively, we superimposed two APC conformations taken from an ensemble of 40 structures generated by the ANM (figure 1A). Measurement of pairwise RMSD between similar residues in these two conformations demonstrated that the RMSD was between 0.15 Angstrom for Gla-domain residues used for matching and 10-15 Angstroms for residues in the catalytic domain. The RMSD could reach 28 Angstroms for some residues in the flexible C-terminal peptides.

**Figure 1.**
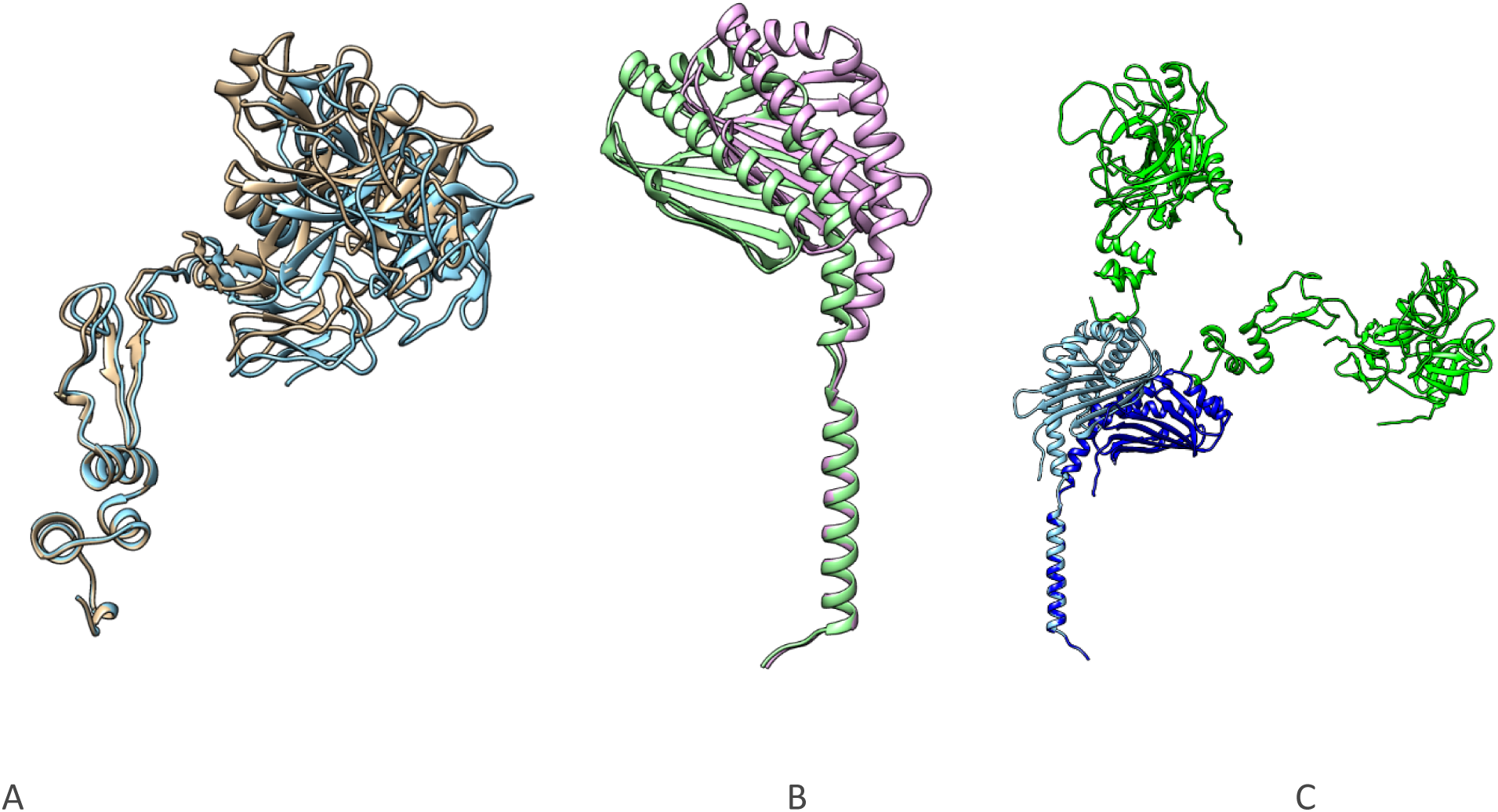
Structures of APC (A) and EPCR (B) that correspond to the full-length proteins, and two representative conformations of the EPCR-APC complex. **A.** Two conformations of APC, derived from ANM analysis (colored in gray and cyan) overlapped with the fixed Gla-domain to demonstrate the flexibility of the APC light chain. Calcium ions in the Gla-domain and other posttranslational modifications are not shown. **B.** The EPCR flexibility according to ANM analysis of the full-length structure. Two overlapping conformations are shown in different colors. Dynamics of EPCR showing the change of the relative positions of the extracellular and transmembrane domains. In addition to hinge-like motions, the extracellular domain of EPCR undergoes rotating motion around the linker sequence, which can change the orientation of the beta-sheet basis of EPCR from parallel to the membrane surface to perpendicular. **C.** APC (green) bound to two different conformations of EPCR (blue and cyan). The representative conformations demonstrate the ability of the EPCR-APC complex to regulate the APC active site distance from the plasma membrane.

### 1.2. EPCR structure and dynamics

EPCR is a type 1 transmembrane protein that exhibits three-dimensional structural homology with the major histocompatibility class (MHC) CD1d. Like the CD1 family of proteins, EPCR has a tightly bound phospholipid in the antigen-presenting groove, and the extraction of lipids results in the loss of PC/APC binding (44). The experimental structure of soluble EPCR bound to the APC Gla-domain has been published (PDB ID: 1lqv.pdb), allowing us to build an accurate structure for the full-length EPCR. The atomic coordinates for the transmembrane domain (TMD) of EPCR missing in the experimental model were taken from the AlphaFold database (model ID: AF-Q9UNN8-F1-model_v2.pdb). The extracellular part of the EPCR structure contains two parallel alpha-helices on top of six beta-sheet platform (figure 1B). A comprehensive analysis of the lipids, found in the EPCR groove, has been reported (45). Lipids bound to EPCR extracellular domain can modulate APC binding, and some lipids have been shown to prevent APC binding. This may prevent APC-EPCR-dependent signaling. Phospholipid removal resulted in the rapid collapse of the hydrophobic cleft of EPCR in our MD experiments that is in agreement with previous publications (46). This collapse changes the structure of ligand-binding regions of EPCR. It suggests that phospholipid supports the functionally active structure of EPCR, although phosphatidylcholine could be exchanged with lysophosphatidylcholine and platelet-activating factor. Replacing phosphatidylcholine by the exchange to these lipids in the hydrophobic groove impaired the ability of EPCR to bind Protein C (47). One report suggests that EPCR can participate in the presenting autoantigens in antiphospholipid syndrome (48).

There is a flexible linking region between the extracellular and transmembrane domains of EPCR. This region also includes the short flexible N-terminal peptide that partially limits large amplitude motions of the extracellular domain of EPCR relative to the TMD and membrane bilayer. The motions of EPCR relative to the plasma membrane are mostly hinge-like motions that change the angle between the extracellular domain of EPCR and the plasma membrane. The RMSD values for two conformations of EPCR selected from the ensemble of 40 conformations generated by the ANM analysis were between 0.172 Angstrom (an exact match) for transmembrane domain residues used for superposition of two structures and 10-18 Angstroms for extracellular domain residues distant from TMD (figure 1B).

### 1.3. Structure of the EPCR-APC complex

It has been demonstrated that protein C, APC, and coagulation factor VIIa (FVIIa) bind to EPCR with high affinity (7,44,49). The Gla-domains of these proteins are responsible for binding to the same extracellular region on EPCR. The experimental crystal structure of the complex of human EPCR bound to the human protein C Gla-domain has been published previously (44). We used this structure as a template for building a full-length EPCR-APC complex (figure 1C). The interaction details of the EPCR-APC Gla-domain have been described in the original publication. Briefly, the Gla-domain binding conformation of EPCR is supported by the phospholipid-bound in the hydrophobic cleft between two α-helices located above β-pleated sheet of EPCR. Hydrophobic interactions (in particular Leu4 and other amino acids located in the ω-like region of the Gla-domain) contribute the most to binding energy between EPCR and APC Gla-domain. The phospholipid does not make direct contact with APC residues. However, the nature of bound phospholipid has a strong effect on APC binding affinity. The lipid-binding groove of EPCR can accommodate many types of lipids (45) resulting in a lipid-dependent change of APC binding. Multiple γ- carboxyglutamic acid-bound Ca^2+^ ions support binding conformation of the APC Gla-domain, but calcium does not participate directly in the EPCR-APC binding (44). The flexibility of the APC light chain between EGF-like domains 1 and 2 allows changes in both the distance from the membrane to the APC active site and the angle between the heavy chain and light chain distal fragment. Combined with the flexibility of the EPCR molecule, the EPCR-APC complex can regulate its conformation and adapt to different binding partners, by adjusting to a wide range of distances from the membrane surface (figure 1C).

### 1.4. Structure of Cav1 oligomer

The structure of the Cav1 oligomer was downloaded from the PDB database (PDB ID: 7sc0.pdb) (50, 51). The Cav1 oligomer described in 7sc0.pdb does not contain the Cav1 fragment known to be located in the cytosol when the Cav1 oligomer is inserted in the caveolar membrane. We used this structure without building the missing part because the topology of the membrane inserted EPCR does not suggest interaction involving the cytosolic part of Cav1. We cannot exclude the interaction of PAR1 C-terminus with the cytosolic part of Cav1 in the intact caveolae, but after initiation of signaling Cav1-PAR1 interaction is no longer detected (2). The structure of Cav1 is shown in figure 2 along with its complexes with the EPCR and PAR1. APC signaling and endothelial barrier protection effects were abolished in cells lacking Cav1, whereas thrombin signaling remained intact (17). This underlines the importance of the Cav1 interactions with PAR1 and EPCR for biased signaling.

**Figure 2.**
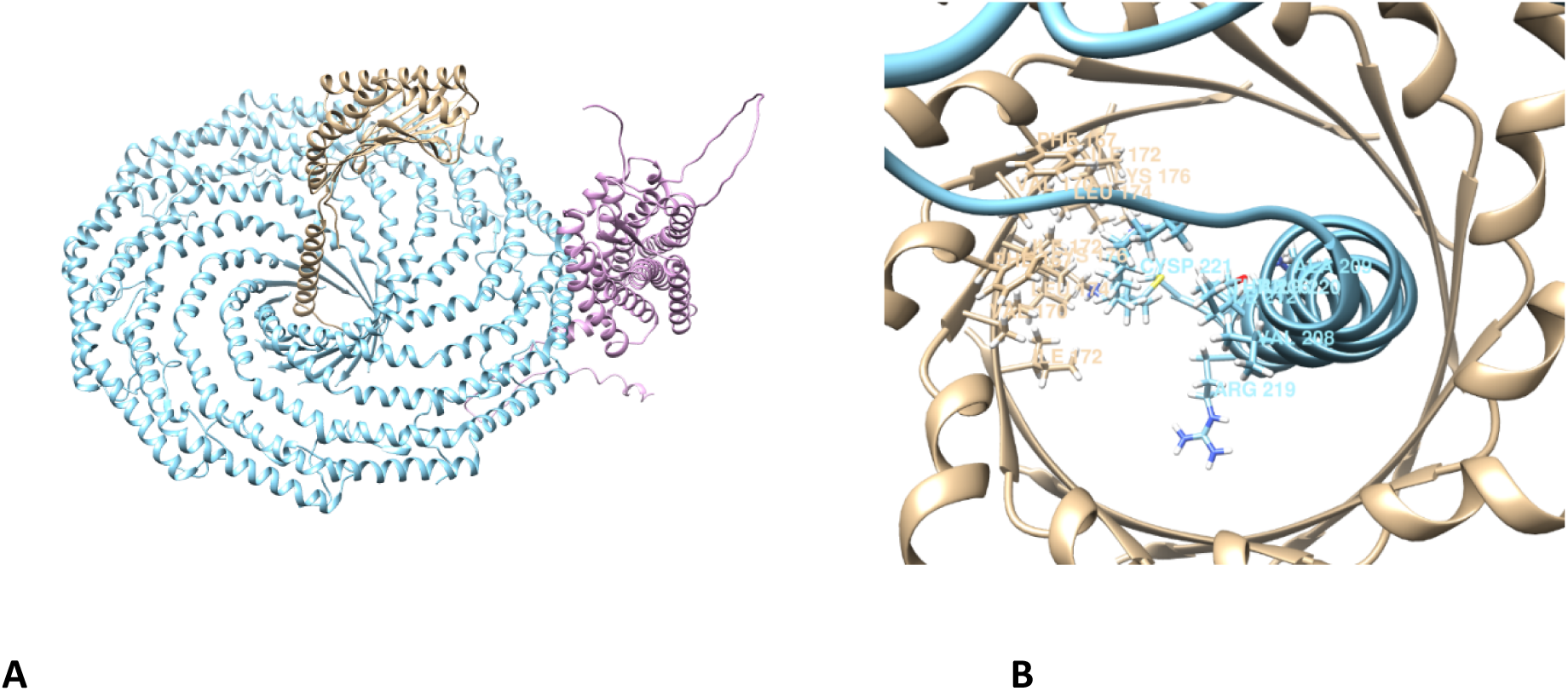
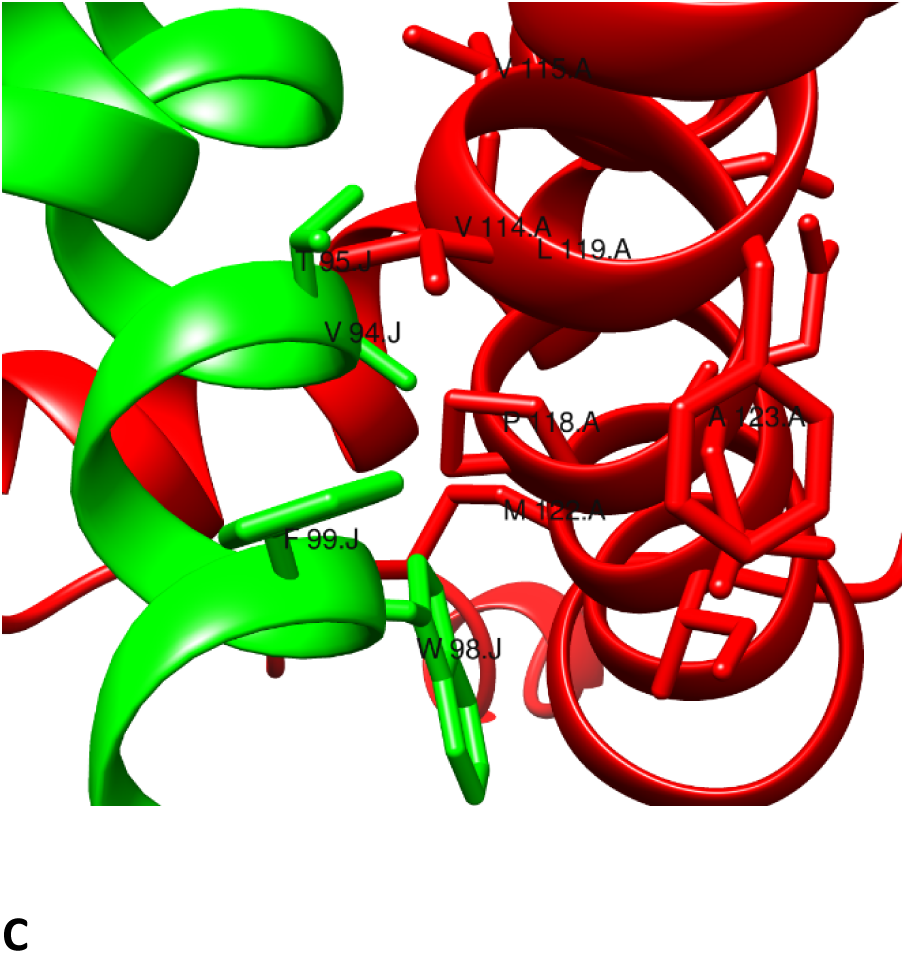
Structures of Cav1-EPCR and Cav1-PAR1 complexes. **A** – overview of the Cav1 oligomer (cyan) bound to EPCR (gray) and PAR1 (magenta). **B** – palmitoylated Cys 221 and transmembrane domain of EPCR (cyan) in the pore-like structure formed by the Cav1 oligomer fragments (gray) showing that palmitoylated EPCR is accommodated by the pore-like structure. Cav1 residues within 5 angstroms from Cys221 are labeled in gray color, while the EPCR residues are labeled in cyan. View from above the surface of Cav1. **C** – Cav1 residues (green color, chain label J) within a 5-angstrom distance of PAR1 residues (red color, chain label A) that belong to TM7. View from above the surface of Cav1 and from the side that allows seeing interacting residues. The red helix in the background represents helix 8 of PAR1.

### 1.5. Structure of PAR1

The experimental structure of PAR1 inhibited by the Vorapaxar has been published (52) and is available for download (PDB ID: 3VW7.pdb). PAR1 is composed of seven transmembrane domains (TM1–7), an extracellular N-terminal domain including 21 amino acid residues encompassing the signal peptide (removed in the expressed PAR1), a domain composed of 20 residues (22-ARTRARRPESKATNATLDPR-41) cleaved by the thrombin and sometimes called propeptide of PAR1, the tethered ligand (57 amino acid residues long), three extracellular loops (ECL1–3), three cytoplasmic intracellular loops (ICL1–3), and an intracellular C-terminal domain (helix 8 of 51 residues) (figure 2). The full-length structure of PAR1 was created by the superposition of the experimental structure of PAR1 with the full- length model downloaded from the AlphaFold database (ID: AF-P25116-F1-model_v3.pdb). The published experimental structure of PAR1 contains a conserved seven TM core structure similar to the δ-subfamily of the class A GPCR to which PAR1 belongs. This structure does not contain the flexible N-terminal peptide of PAR1 which is cleaved by thrombin, APC, or other proteases to form a TL that initiates PAR1 signaling. The experimental structure also does not contain the C-terminal domain of PAR1 (52). Atomic coordinates for these fragments were taken from the AF-P25116-F1-model_v3.pdb and added to the full-length structure of PAR1. The structure of PAR1 is shown in figure 2 in the complex with Cav1, and also in figures 4 and 6 in the signaling complexes.

### 1.6. Structure of β-arrestin-2

The experimental structure of human β-arrestin-2 is not known, however, the homologous structure of the rat (53) (PDB ID: 6K3F) β-arrestin-2 and structures of other members of arrestin family including structurally similar PAR1 complexes of GPCRs with arrestins [i.e., rhodopsin-arrestin complex (54)] have been published. In this work, we used protein containing 409 amino acid residues (UniProt ID: P32121 ARRB2_HUMAN) and the model of human β-arrestin-2 (ID: AF-P32121-F1-model_v3.pdb) to create models of β-arrestin-2 and its complexes with PAR1 or PAR1-EPCR (figure 6). AlphaFold produces a per-residue confidence score that was < 50 for some loops and the C-terminus of β-arrestin-2, suggesting that these regions are flexible and may be unstructured.

### 1.7. Interactions of the PAR1 N-terminal peptide with the APC-EPCR complex

#### 1.7.1 Interactions in the catalytic domain of APC

The details of the binding of the PAR1 N-terminus to APC-EPCR are incompletely understood. APC cleaves PAR1 at R46, while both thrombin and APC can cleave PAR1 at R41 (55). We used homology modeling, flexible peptide-protein docking, and MD simulation of the system composed of APC and PAR1 N-terminal peptide to gain insights into APC-PAR1 interactions in the situations leading to either R41 or R46 cleavage. To build the structure of the APC catalytic domain interacting with the PAR1 N-terminus we used a homology modeling method that employed an experimental structure of thrombin in complex with uncleaved PAR1 (PDB ID: 3lu9.pdb) as a template. Interactions taking place in the catalytic cleft of APC during cleavage at R41 should be similar to the interactions in the thrombin-PAR1 peptide complex resulting in the same R41 cleavage (56). This model is presented in figure 3A, which shows the catalytic cleft of APC with the PAR1 R41 in a position enabling its hydrolysis by the catalytic triad of APC. We modeled PAR1 cleavage by APC at R46 using the R41 cleavage model as a template with two strategies. In the first approach, the model of PAR1 N-terminus containing residues 33-57 was extended to include residues 22-32, and the resulting peptide was moved to match the position of R46 with the position which has been occupied by the R41 in the template. After removing the steric clashes using the structure editing tool of UCSF Chimera, this structure was simulated with MD. In the second approach, to build the structure of the APC-PAR1 complex that is suitable for the cleavage at R46, we employed site-directed mutagenesis to replace the PAR1 sequence 33-57 with the sequence 38- 57 (after this procedure R41 became R46) (figure 3B). Possible amino acid side chain clashes were removed by the structure editing tool which selected the correct rotamers. Using the same tool, the 38-57 sequence was extended to build the structure of the complete PAR1 N- terminus (22–98) with the R46 in the active site of APC in the proper position for cleavage. This structure was also studied by the MD simulation and following analysis of the MD trajectory.

**Figure 3.**
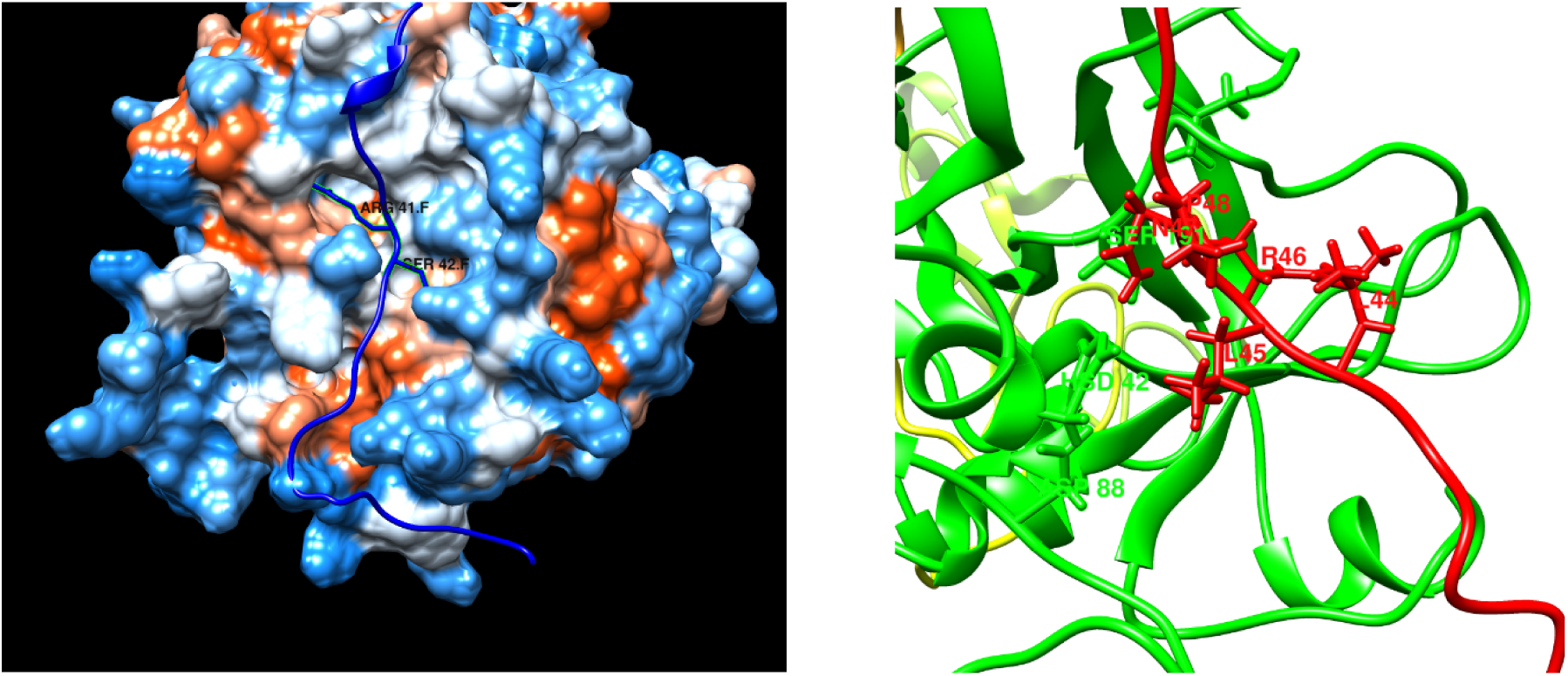
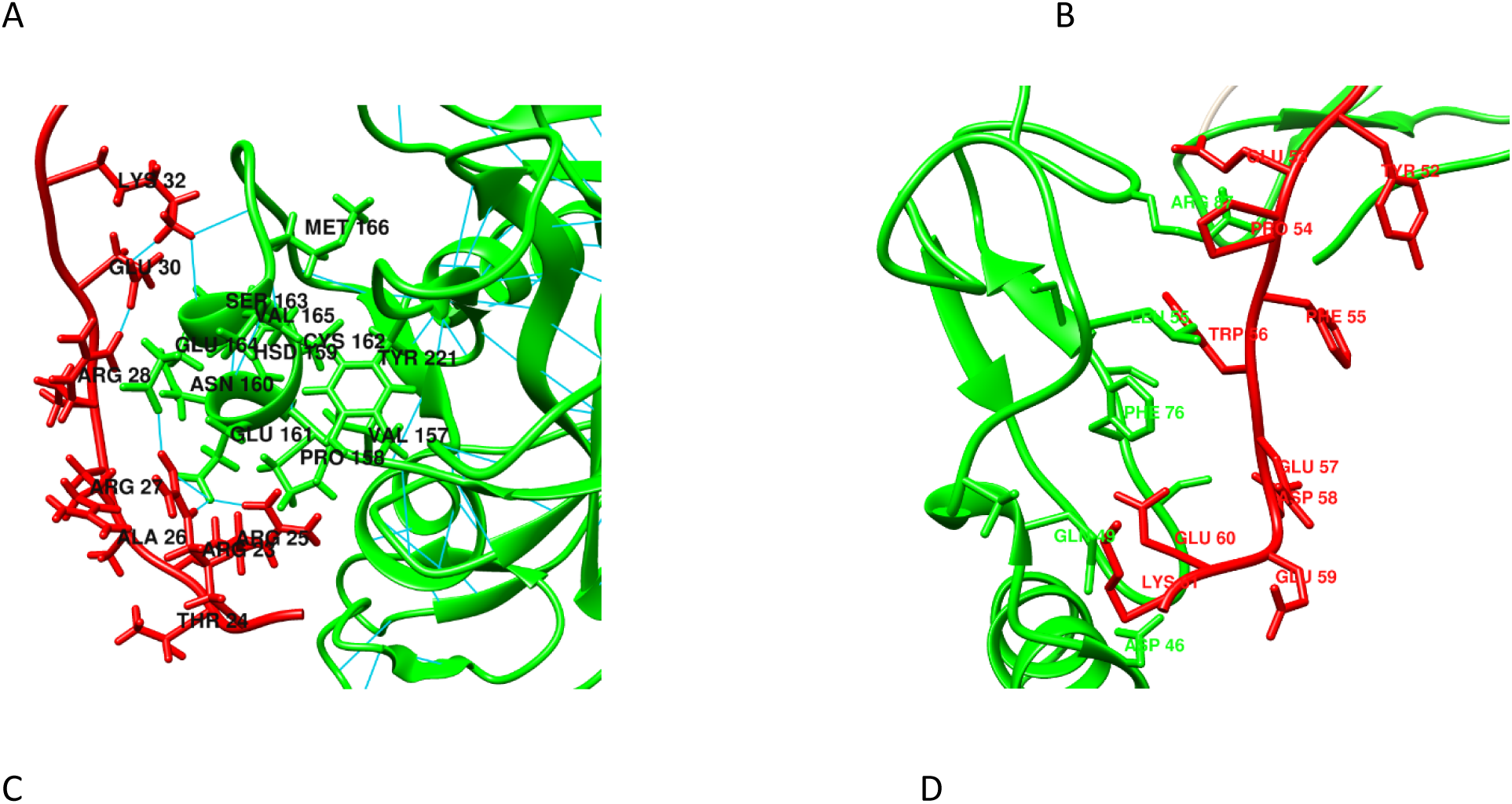
Interaction of the PAR1 N-terminal peptide with the catalytic domain (panels A, B, and C) and light chain of APC. **A**. Hydrophobicity surface representation of the PAR1 peptide in a position appropriate for the R41 cleavage. Side chains of R41 and S42 are shown as sticks in blue color. **B**. PAR1 peptide (red) in the active site of APC (green) in a position that should result in cleavage at R46. PAR1 residues 44-48 are labeled, and their side chains are shown. Similarly, APC catalytic triad residues HIS, ASP, and SER residue side chains are shown and labeled as HSD 42, ASP 88, and Ser 191. These labels correspond to the active site triad His57, Asp102, and Ser195 in the chymotrypsinogen numbering, respectively. **C.** Interaction of the PAR1 N- terminus (red) with the helix 162 residues of APC (green) in the peptide position related to the R46 cleavage. The peptide residues can form numerous contacts with the 162 helix residues and have a longer residence time in the MD simulation as compared to the simulation of peptide in the R41 cleavage position. The Glu170 and Glu167 (chymotrypsinogen numbering) correspond to the Glu 164 and Glu 161 labels in the figure, respectively. **D**. Interaction of the tethered ligand (red) with the APC light chain (green). The amino acid residues of APC EGF1 domain in a distance less than 5 Angstrom from PAR1 N-terminal peptide are labeled with residue name followed by the residue number in a color corresponding to the chain to which the residue belongs.

Two residues unique for the 162-helix of APC on the catalytic domain (Glu-167 and Glu-170), constitute a specific PAR-1-binding exosite on APC that facilitates the recognition and cleavage of the receptor by the protease on the endothelial cell surface (57). The model of the signaling complex suggests that Glu-167 and Glu-170 residues may interact with the PAR1 N-terminal peptide (22-ARTRARRPESK-32) representing a part of the biologically active peptide (58) removed from PAR1 by the thrombin or APC cleavage. MD simulation of the system composed of structure derived from the homology model of APC active site – PAR1 peptide in the position proper for R41 cleavage allowed to demonstrate short-lived hydrogen bond formation between Arg-23 of PAR1 and Glu-167 of APC (not shown). The simulation of the PAR1 peptide in the APC catalytic cleft in a position suitable for R46 cleavage demonstrated that the 22-37 peptide (with additional 5 amino acid residues as compared to the R41 cleavage peptide) makes more contacts with the 162 helix and binds to Glu-167 and Glu-170 region (figure 3C) with longer residence time as compared to R41 cleavage complex. This suggests tighter binding of PAR1 N- terminus to APC catalytic domain in the position preferable for R46 cleavage. It is interesting to compare this complex with the PAR1 cleavage by thrombin when tight binding of the hirudin-like sequence of PAR1 to the thrombin exosite positions R41 at the active site of thrombin for cleavage. This observation may at least in part, explain the preferential cleavage of PAR1 by APC at the R46 position.

#### 1.7.2. Interactions of TL with the light chain of APC

After cleavage, before tethered ligand binding to PAR1 to activate it, the PAR1 N-terminus interacts with the light chain of APC in two regions: C-terminus including Lys-150 and Lys-151 (59), and EGF-like domain 1 of APC (60). We modeled these interactions using several approaches: in the first approach, the PAR1 N-terminal peptide fragments were docked to APC using the flexible peptide docking method (29); the second approach included folding of PAR1 and EPCR-APC as heterotrimer with AlphaFold software; and the third approach was MD simulation of the system containing APC and PAR1 N-terminus.

Models suggest that simultaneous binding of TL to the active site and interactions with the APC light chain is unlikely due to geometric limitations. This suggests a sequence of events including cleavage at R41 (or at R46) resulting in the formation of TL, dissociation of TL from the APC catalytic domain, and interaction with the APC light chain and then TL PAR1 interactions leading to the PAR1 activation. Docking and MD simulation experiments show that interactions of the PAR1 TL peptide with the APC light chain are transient and can involve different fragments of the peptide. A model of one of the binding scenarios is shown in figure 3D. Most likely, these are low-affinity interactions, because interacting regions did not stay in contact for the entire time of the MD simulation. Considering that the residence time is reciprocal of the off rate, the binding affinity of these fragments is expected to be low. Most often, the PAR1 sequence 58- DEEKNESGLTEYRLVS-72 was found in contact with the APC EGF-like 1 domain, suggesting that part of the hirudin-like sequence is involved in binding. The peptide fragments closer to the N- terminus of TL were found forming transient contacts with the light chain C-terminus (including K150 and K 151), which was flexible (RMSD about 28 Angstrom) and was located close to the EGF-like 1 domain of APC.

## 2. Structure of the complexes EPCR-Cav1, EPCR-PAR1, and Cav1-PAR1

### 2.1. EPCR-Cav1 complex

The EPCR cytoplasmic fragment contains only three amino acids (Arg-Arg-Cys) (61), and it was reported that Cys residue could be palmitoylated (62). Because palmitoylation is reversible (63), we tested palmitoylated and intact EPCR interaction with the PAR1. The very short cytoplasmic tail of EPCR and its possible palmitoylation suggests that EPCR cytosolic tail will be localized at the level of polar heads of lipids in the inner leaflet of the membrane bilayer when palmitoylated, or very close to the membrane surface without palmitoylation. To identify the binding site for EPCR on the Cav1 oligomer, we performed docking experiments with both, intact EPCR molecule and its TMD fragments using the HDOCK server. According to the cryo-electron microscopic data, the Cav1 oligomer complex is composed of 11 protomers organized into a tightly packed disc with a flat membrane-embedded hydrophobic surface with a pore-like structure in the center capable of accommodating residues from another protein (51). An analysis of docking results demonstrated that there are two types of binding sites for EPCR TMD on the cav1 oligomer. The first type of binding site is located in the pore-like structure of Cav1 and this site can accommodate both, palmitoylated and intact C-termini of EPCR. As shown in Figures 2A and 2B, the C-terminal residues of EPCR are located in the pore-like structure formed by the C-terminal fragments of Cav1 monomers. Amino acid residues participating in the binding were: 215-CTGGRRC-221 from EPCR C-terminus and 166-IFSNVRINLQKE-177 at the Cav1 C-terminus. The second type of binding site was identified in folding experiments and was located at the rim of Cav1 (not shown). The binding site involves two protomers and includes amino acid residues 103-107 of one protomer in addition to the residues 83-87 of the second protomer that interacts with the EPCR TMD fragment (residues 204-213) close to the C-terminus. Because there are 11 binding sites per one Cav1 oligomer, and these sites are readily accessible as compared to the central pore-like structure, we speculate that the binding sites at the rim of Cav1 are more functionally important. It is also highly likely that EPCR can bind to the Cav1 bound PAR1 without direct contact with the Cav1, like an EPCR bound to PAR1 in figures 4B and 6A.

**Figure 4.**
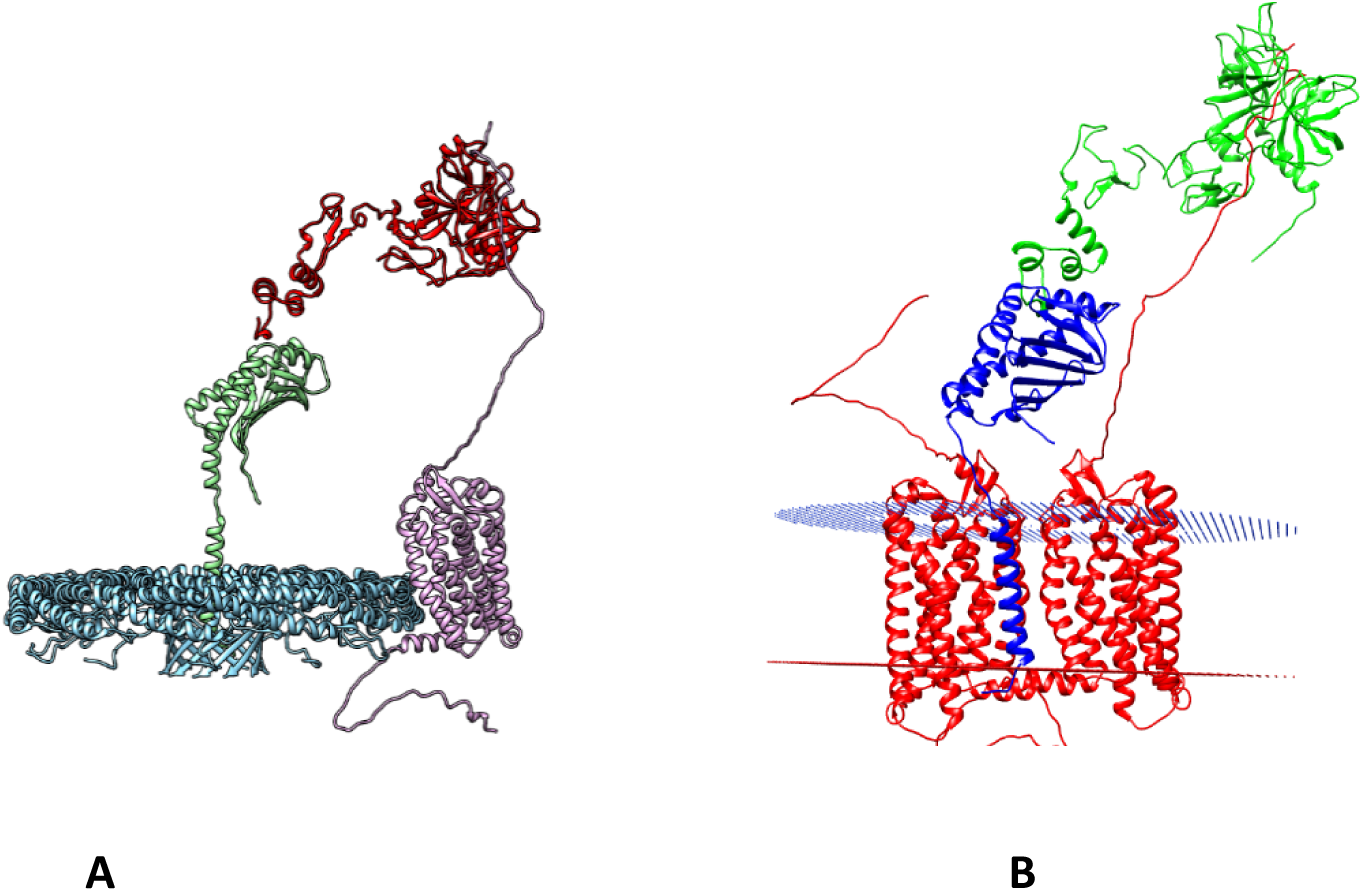
Hypothetical APC induced PAR1-biased signaling complexes before formation of the tethered ligand. **A**. Cav1-EPCR-APC-PAR1 complex. **B**. complex of EPCR-APC with PAR1 dimer representing possible model of PAR1 transactivation. The plasma membrane is shown using dummy atoms placed in positions of phospholipid polar heads.

### 2.2. Building full-length three-dimensional structures of EPCR-PAR1, and EPCR-PAR1-PAR1 ternary complex

EPCR and PAR1 can be co-isolated by immunoprecipitation of cell homogenates and detected by Western blotting suggesting these proteins are co-localized and can interact with each other directly or indirectly (17). It is also known that PAR1 can form functionally active homodimers or heterodimers with PAR2 (1, 4). The binding sites involved in the formation of PAR1 oligomers are not known, it is possible that the dimerization of different PARs involves different regions. It has been demonstrated that the TM4 helix of PAR4 participates in dimerization (64). The specific structural rearrangements that occur following activation by the tethered ligand and how cofactors and heterodimers influence the receptor allostery are all open questions (4). To establish regions of PAR1 involved in dimerization and possible interactions of EPCR TMD with PAR1, we created EPCR- PAR1, PAR1-PAR1, and ternary EPCR-PAR1-PAR1 complexes using the AlphaFold method. This method is the most accurate method for creating protein models from their sequences as well as creating homo- or hetero-oligomeric protein complexes (34, 38). Predicted models of the EPCR-PAR1 complex demonstrated two binding modes of EPCR TMD to PAR1. The first binding mode places EPCR TMD at the interface formed by transmembrane helices 4 and 5 of PAR1. The second binding mode involves the EPCR TMD contacting helices 6,7, and cytoplasmic helix 8 of PAR1 (figure 6A). The region of EPCR that links extracellular and transmembrane domains was close to the PAR1 region connecting the transmembrane helix 1 and the N-terminal peptide in this mode. We noted that the outcome of the AlphaFold method for building hetero-oligomeric complexes depended on the order of protein sequences at the input stage. Therefore, PAR1 complexes were prepared using several AlphaFold runs while changing the order of sequence input. The complexes for further analysis were selected from the highest-scoring complexes after applying all known constraints derived from the membrane topology of these proteins and known experimental observations. Selected models representing an EPCR-PAR1 model, and a signaling complex composed of EPCR-APC-PAR1 were further investigated with MD simulations. The selected complex included the interaction of the EPCR TMD with the TM6, d TM7, and helix 8 of PAR1.

### 2.3. Cav1-PAR1 complex

As in the case of EPCR and PAR1 interaction, Cav1 and PAR1 can be co-isolated by immunoprecipitation before initiation of PAR1 signaling, suggesting direct or indirect binding of PAR1 to Cav1 in the caveolae (2,17,18). A peptide related to the scaffolding domain (CSD) of Cav1 can bind to alpha subunits of G proteins, including those involved in PAR1 canonical signaling (65). Moreover, CSD binds some G protein coupled receptors (GPCRs) (66). We hypothesize that CSD and some other amino acid residues of Cav1 participate in the Cav1-PAR1 complex (figure 2A) formation and build the complex in which relative orientations of PAR1 and Cav1 correspond to the topology of the complex in the membrane bilayer. First, we docked Cav1 CSD to PAR1 structure and selected resulting structures that are consistent with the membrane topology of the Cav1 and PAR1. Then we superimposed PAR1-bound CSD with the CSD in the Cav1 oligomer structure. Next, we inspected superimposed structures and selected the ones that satisfy the topological requirements. Figure 2C shows an example of the selected structure in which PAR1 helix 8 is directed toward the pore-like structure of Cav1. Another example of a complex that satisfies the topological requirements had a CSD binding site located near residues 201-221 of PAR1 which are related to the cytosolic fragments of TM3 and TM4 and ICL connecting these two domains.

## 3. The model of APC-mediated PAR1 biased signaling complex

The biased PAR1 signaling by APC or thrombin involves cleavage of the PAR1 N-terminus at either Arg41 or Arg46 (2,16,18). It has been hypothesized that biased PAR1 signaling involves the interaction of the receptor with the first EGF-like domain of APC (60). Based on the literature, a model of APC-biased PAR1 signaling should describe the role of Cav1, which interacts with EPCR and PAR1 at the initial stages of signaling. The model should also provide the structural basis for the interaction of the EPCR-APC complex with PAR1, and interactions of the activated PAR1 with the b-arrestin-2. In this work, we modeled the initial stages of PAR1 signaling including the formation of the PAR1-b-arrestin-2 complex.

At least three variants of initial signaling complex structure that can support PAR1 N-terminal peptide cleavage by APC and formation of the tethered ligand can be deduced from the published observations and structural models. In the first variant of the PAR1 cleavage complex APC-EPCR is bound to the pore-like region, while PAR1 is bound to the rim of the Cav1 oligomer (figure 4 A). Cav1 can accommodate the PAR1 cleavage complex in which both PAR1 and EPCR are bound to the rim of Cav1 (not shown). According to the experimental observations, binding of APC to EPCR induces dissociation of EPCR from Cav1 (18), therefore this complex most likely is a transient complex with a short lifetime. The geometry of this quaternary complex places the APC catalytic domain and PAR1 at an optimal distance to enable the interaction of extended conformation of the PAR1 N-terminus with the active site of APC, leading to its cleavage and formation of the TL.

The second variant of the signaling complex can form in the lipid rafts without Cav1 oligomer in which case the interactions between APC, EPCR, and PAR1 can be similar to the first type of signaling complex (not shown). The sequence of events leading to the formation of this complex is not known. One could speculate that this is the same structure as in the first complex after dissociation of Cav1 and translocation of APC-EPCR-PAR1 to the lipid raft. It is also equally likely that this is a newly formed complex. Like the type one complex, a certain distance between PAR1 and EPCR-APC is required for this signaling complex to optimally bind and cleave the PAR1 N-terminus to form the TL.

The third variant of the signaling complex is composed of EPCR-APC bound to the PAR1-PAR1 dimer (figure 4 B). The existence of this complex is supported by the well-documented PAR1 homodimers and PAR1-PAR2 heterodimers that support PAR1 signaling by the transactivation mechanism and by dimerization of other PAR receptors (1,16,64). This complex can have two subtypes. The first is composed of EPCR-APC bound to PAR1 dimer only by the PAR1 N-terminus of one of the protomers (not shown), similar to the second type described in figure 4 B. The second subtype includes two simultaneous links; EPCR TMD directly bound to PAR1 TM helices and the binding of the PAR1-N-terminal peptide to the APC catalytic domain (figure 4 B).

Experimental evidence for the interaction of EPCR with PAR1 has been provided by the co-immunoprecipitation of these proteins from cell lysates (3,17,67). These experiments did not identify interaction sites and did not exclude the indirect binding of PAR1 to EPCR via a third participant. Our modeling experiments, especially EPCR-PAR1 heterodimers generated with the AlphaFold2 suggest that binding between EPCR and PAR1 most likely is mediated by the interaction of the transmembrane domains of these proteins.

The original publication on the experimental three-dimensional structure of PAR1 used MD simulations to investigate structural changes related to the activation of PAR1, demonstrating that MD simulations can be informative in the study of PAR1 activation (52). Therefore, we used MD simulations to investigate the structure and dynamics of PAR1 in the signaling complex at the atomic level. First, we investigated the interaction of the R41 or R46 cleaved TL with the ternary PAR1-EPCR-β-arrestin-2 complex by MD simulation of the models.

Previous molecular modeling studies suggested a shallow ligand binding site for R41 cleaved TL located on the surface of the receptor PAR1 (52). Mapping the binding site on PAR1 for its agonist peptide SFLLRNP identified four PAR1 surface residues Leu96, Asp256, Glu260, and Glu347 contacting the peptide (69). It was demonstrated that Phe2 and Arg5 are peptide activity-determining residues with an edge-to-face CH/π interaction, mediated by the Phe2 of the SFLLRNP peptide, that is essential for the aggregation of human platelets via PAR-1 activation (70). Our MD simulation of the PAR1-EPCR complex with EPCR TMD interacting directly with the PAR1 TM helices 6 and 7 demonstrated that EPCR facilitates the binding of Phe43 with the surface residues of a hydrophobic pocket of PAR1. This is the same hydrophobic pocket where PAR1 antagonist Vorapaxar binds (52). Consistent with the edge-to-face CH/π interaction hypothesis of PAR1 activation, Phe43 in the inserted ligand is positioned to form these interactions with Tyr residues (figure 5C). The activity determinants for the R46 cleaved TL are not known and our docking experiments did not reveal edge-to-face interactions for this peptide. To characterize an interaction of R46 TL with the PAR1, we provide one of the frames taken from the MD simulation trajectory (figure 5D) showing interactions between R46 cleaved TL and PAR1 surface residues during PAR1 activation.

**Figure 5.**
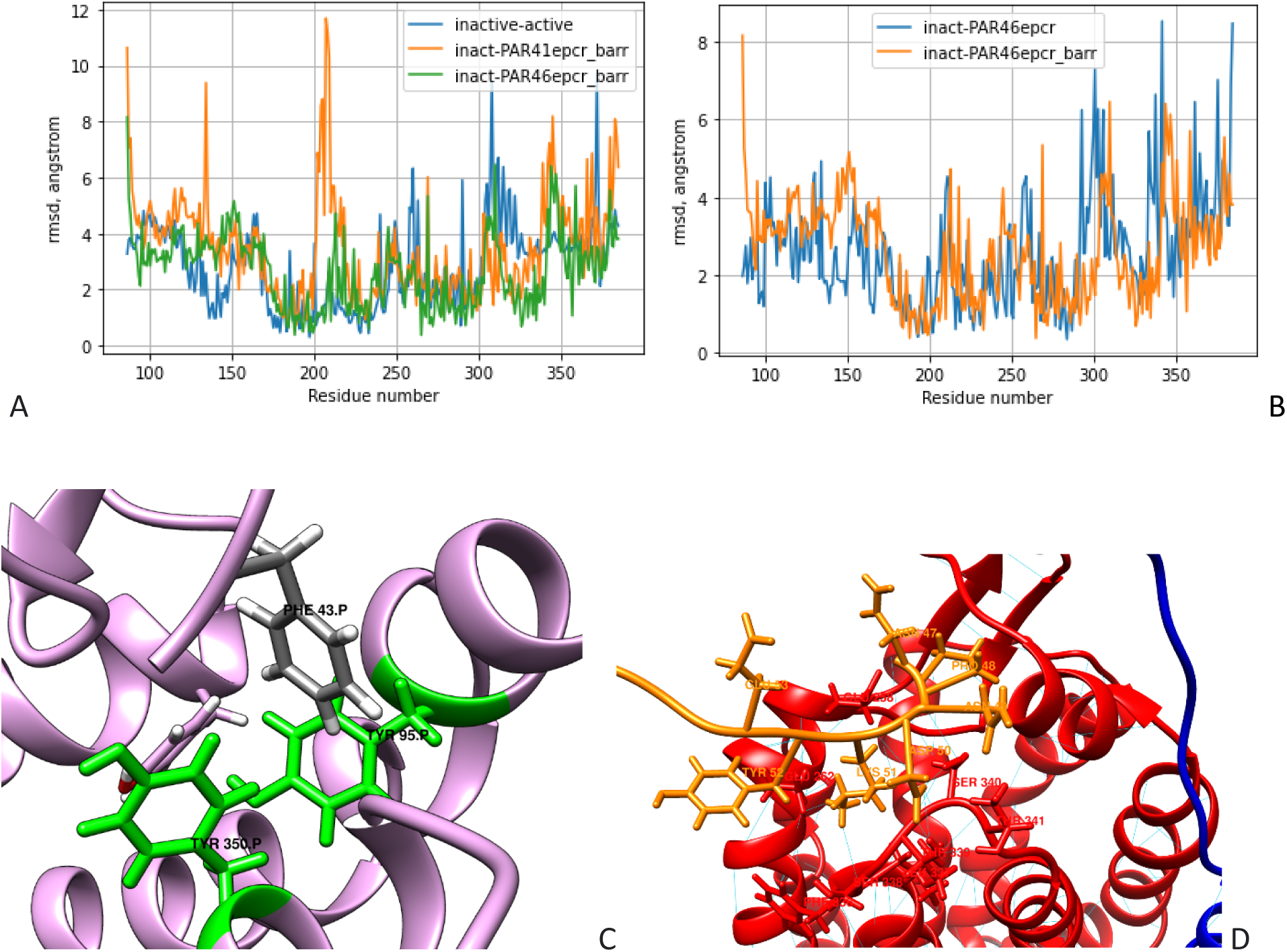
Models of PAR1 activation by TL in APC-biased PAR1 signaling. **A**. Pairwise RMSDs for the same residues in the overlapped active and inactive PAR1 molecules downloaded from the GPCR database (shown in blue and labeled inactive-active). Pairwise RMSDs between PAR1 (A86-S396) residues before and after MD simulation of the PAR1-EPCR-β-arrestin-2 complex with the PAR1 TL cleaved at the R41 (orange, labeled inact-PAR41epcr_barr). Pairwise RMSDs between PAR1 (A86-S396) residues before and after MD simulation of the PAR1-EPCR-β-arrestin-2 complex with the PAR1 TL cleaved at the R46 (green, labeled inact- PAR46epcr_barr). **B.** Pairwise RMSDs of PAR1 cleaved at R46 in complex with EPCR as compared to the PAR1-EPCR-β-arrestin-2 complex. The pairwise RMSDs for PAR1-EPCR complex before and after MD simulation (blue, labeled inact-PAR46epcr). The pairwise RMSDs for PAR1-EPCR-β-arrestin-2 complex before and after MD simulation (orange, labeled inact- PAR46epcr_barr). **C**. Edge-to-face interaction of Phe43 (metallic color) with aromatic (Tyr) amino acids (colored green) involved in the PAR1 activation by the TL cleaved at R41. **D**. Interaction of the R46 cleaved TL (orange) with the extracellular region of PAR1 (red). The blue structure is a small fragment of the EPCR backbone. The interactions of the TL after 50 ns MD simulation of the PAR1-EPCR complex are shown.

It has been shown that activation of PAR1 results in small changes in the relative positions of TM helices, the intracellular and extracellular loops, and the side chains of amino acid residues (52). Because these multiple subtle changes make it difficult to differentiate between active and inactive PAR1, we used active and inactive conformations of PAR1 downloaded from the GPCR database (71, 72) as reference structures. The goal of the experiments described in Figures 5A and 5B is to follow the conformational changes related to PAR1 activation by R41 and R46 cleavage in the presence of EPCR and β-arrestin-2. To achieve this goal, we measured pairwise RMSD between the same residues in the active and inactive PAR1 reference structures and in the PAR1 complexes before and after MD simulations. However, there is a caveat: the PAR1 conformations in the AlphaFold models of complexes used for MD simulations did not match either active or inactive reference structures exactly. Therefore, the results described below should be considered as PAR1 adaptation to the conditions of biased signaling, rather than conformational changes related to the inactive to active PAR1 conversion. Because PAR1 N-terminus and C-terminus are unstructured, the receptor alignment and superposition were based on the body of PAR1, on its transmembrane domains. We compared RMSDs for the reference structures (inactive - active) with the RMSDs for PAR1 in signaling complexes before (inactive) and after MD simulation (active), with the understanding of the caveat described above. To simplify the comparisons, we used RMSD equal to 4 Angstrom as a threshold value and considered RMSD larger than the threshold during the comparisons. The regions of reference PAR1 structures with large conformational changes include neighboring residues for residue numbers: 165, 240, 260, 285, 315, and 380, which correspond to ECL1, ECL2, ECL2, TM5, TM6, and helix8, respectively (figure 5A blue line). Pairwise RMSDs between PAR1 residues before and after MD simulation of the PAR1-EPCR-β-arrestin-2 complex with the PAR1 TL cleaved at the R41 (figure 5A orange line) demonstrated conformational changes in the following regions.

Residue numbers 100-140, 200-220, around 270, around 310, and 340-370, corresponding to the PAR1 regions TM1 with ICL1, ICL2, TM5, TM6, and TM7. MD simulation of the signaling complex PAR1-EPCR-β-arrestin-2 with the PAR1 TL cleaved at the R46 (figure 5A green line) demonstrated changes: at 140-150, around 220, 270, 300-320, and 340-350 that belongs to PAR1 regions TM2, TM4, TM5, ICL3-TM6, and ECL3-TM7. A comparison of changes related to PAR1 binding of R41 cleaved TL with the changes related to binding of R46 cleaved TL demonstrated that the changes involve similar regions, although involved residues match only partially. A comparison of RMSD for reference structures (blue line) with the RMSD related to R41 cleaved TL (orange line) and R46 cleaved TL highlighted the changes in the extracellular loops (ECL1-ECL2) for reference structures while changes in the RMSD for intracellular loops (ICL1-ICL3) and adjacent to ICLs regions of transmembrane domains were observed for orange and green lines. In summary, these comparisons reveal that there are many regions of PAR1 that undergo small conformational changes in all three situations described in figure 5A. The residues in the TM5 and TM6 regions undergo RMSD changes of more than 4 Angstrom in all three situations. Overall, these observations are consistent with the existence of multiple conformations (an ensemble) of activated PAR1 (73). Because the presence of EPCR enables the biased signaling irrespective of the TL cleavage site, we compared RMSDs for EPCR-PAR1 complexes with or without β-arrestin-2 to identify EPCR- induced changes in PAR1 related to the biased signaling (figure 5B). The RMSD changes larger than 4 Angstrom involved PAR1 residues 110, 130, 220, 260, 280-320, 330-340, which represent regions TM1, ICL1, TM4, ECL2, TM5-ICL3-TM6, and TM6-ECL3 (figure 5B, blue line) for the EPCR-PAR1 complex. RMSD changes in the presence of β-arrestin-2 include residues 130-150, 220, 240, 275, 310-320, and 350-360 that represent PAR1 regions ICL1-TM2, TM4, TM4, TM5, TM6, and TM7, respectively (figure 5B, orange line). The results demonstrated changes in extracellular loops ECL2 and ECL3 in these two situations. Interestingly, PAR1-EPCR and PAR1-EPCR-β-arrestin-2 complexes undergo conformational changes in the regions that include TM5 and TM6 suggesting that EPCR promotes PAR1 changes that favor β-arrestin-2 binding. The absence of large RMSD changes in the ICL3 in the presence of β-arrestin-2 can be explained by the binding of ICL3 residues to β-arrestin-2 which limits the conformational changes in this loop. These results suggest an important role of EPCR TMD binding to the PAR1 in the initiating of biased signaling.

During building the models of the Cav1-PAR1-β-arrestin-2 complex, we noted that membrane binding C domain edge of β-arrestin-2 may be prevented from membrane binding in some relative orientations of PAR1 and Cav1 oligomer. This was observed for the model where PAR1 TM3 and TM4 formed Cav1 binding interface. In this model, the C domain edge was blocked from the membrane by the surface of the Cav1 oligomer. Because membrane binding is important for G protein function, we created Cav1-PAR1-G protein complex models and analyzed the potential effect of Cav1 on PAR1-G protein function. We found that Cav1 can prevent membrane binding of G proteins in many relative orientations of PAR1-G protein and Cav1. Analysis of the literature data on the activation of G proteins suggests that interaction with the Cav1 may serve as a mechanism that selects β-arrestin-2 as a downstream signaling molecule instead of G proteins. Two factors can contribute to this selection mechanism. First, caveolin binds to G proteins via its CSD (65, 66), and this binding can block G protein function by competing with G protein binding to PAR1. Second, blocking the functionally essential membrane binding of G protein by the Cav1 in some relative orientations of Cav1 and PAR1-G protein. Considering that β-arrestin-2 can be activated without membrane contact of the C edge loops (73–75) the caveolar localization and interaction with Cav1 can provide advantages for the functional activity of the PAR1-β-arrestin-2 complex, but not the PAR1-G protein complex. Additionally, PAR1 cannot couple to the G protein to initiate canonical signaling if the C-terminal cytoplasmic domain of the receptor was phosphorylated by G protein coupled receptor kinases (GRK). Instead, β-arrestin-2 is recruited to bind to the phosphorylated receptor, thereby initiating biased PAR1 signaling. The next section describes the possible role of PAR1-bound EPCR TMD in the β-arrestin-2 biased signaling.

### 4.2. Activation of b-arrestin-2 and interaction of the EPCR C-terminal residues with the PAR1 bound b-arrestin-2

According to published data, there are two major interaction interfaces between PAR1 and β-arrestin-2 (74, 75). The first is a β-arrestin-2 N-domain sequence that binds to the phosphorylated C-terminus of PAR1 and the second is a finger-loop-region (FLR) inserted into the intracellular PAR1 cavity (74, 75). Activation of β-arrestin-2 includes conformational changes in the loop encompassing residues between R66 -S75 of FLR inserted into a pocket formed by the TM5, TM6, ICL3, and TM7 of PAR1. Activation also includes change of the angle between the N and C domains of β-arrestin-2 (74, 75). Additionally, the C-edge loops of arrestins can act as a membrane anchor to stabilize the PAR1-β-arrestin complex (77). Activated β-arrestin-2 can have different conformations, which in part are mediated by phosphorylation of at least six residues including Thr178, Ser194, Ser267, Ser268, Ser281, Ser361 and Thr383 (76). It has been demonstrated that in the APC biased signaling EPCR occupancy recruits GRK5, which phosphorylates PAR1 cytoplasmic C-terminal domain, leading to binding of β-arrestin-2 to PAR1 (2). For the β-arrestin-2 biased signaling, in addition to binding of the β-arrestin-2 N domain to the phosphorylated PAR1 C terminus, the FLR loop needs to bind to the body of PAR1 and block the G protein binding to PAR1 and its subsequent G protein dependent signaling. In the biased signaling model, the EPCR-PAR1-β-arrestin-2 complex formation places EPCR C-terminus in the region where it can interact with the PAR1- bound b-arrestin-2 directly or indirectly (figure 6A). To reveal the effect of EPCR on the β- arrestin-2 activation we generated PAR1-β-arrestin-2 models with AlphaFold software and compared these models to the PAR1-β-arrestin-2-EPCR ternary complex models. We found that the presence of EPCR TMD bound to PAR1 at the interface between TM6 and TM7 did not create hindrance for PAR1-b-arrestin-2 binding. Two types of b-arrestin-2 activation related conformational changes are: the FLR changes - finger like loop is converted into omega-like loop and rotation of N and C domains of b-arrestin-2 relative each other. Both these changes were detected in the MD simulation trajectory of the PAR1-EPCR-b-arrestin-2 complex.

**Figure 6.**
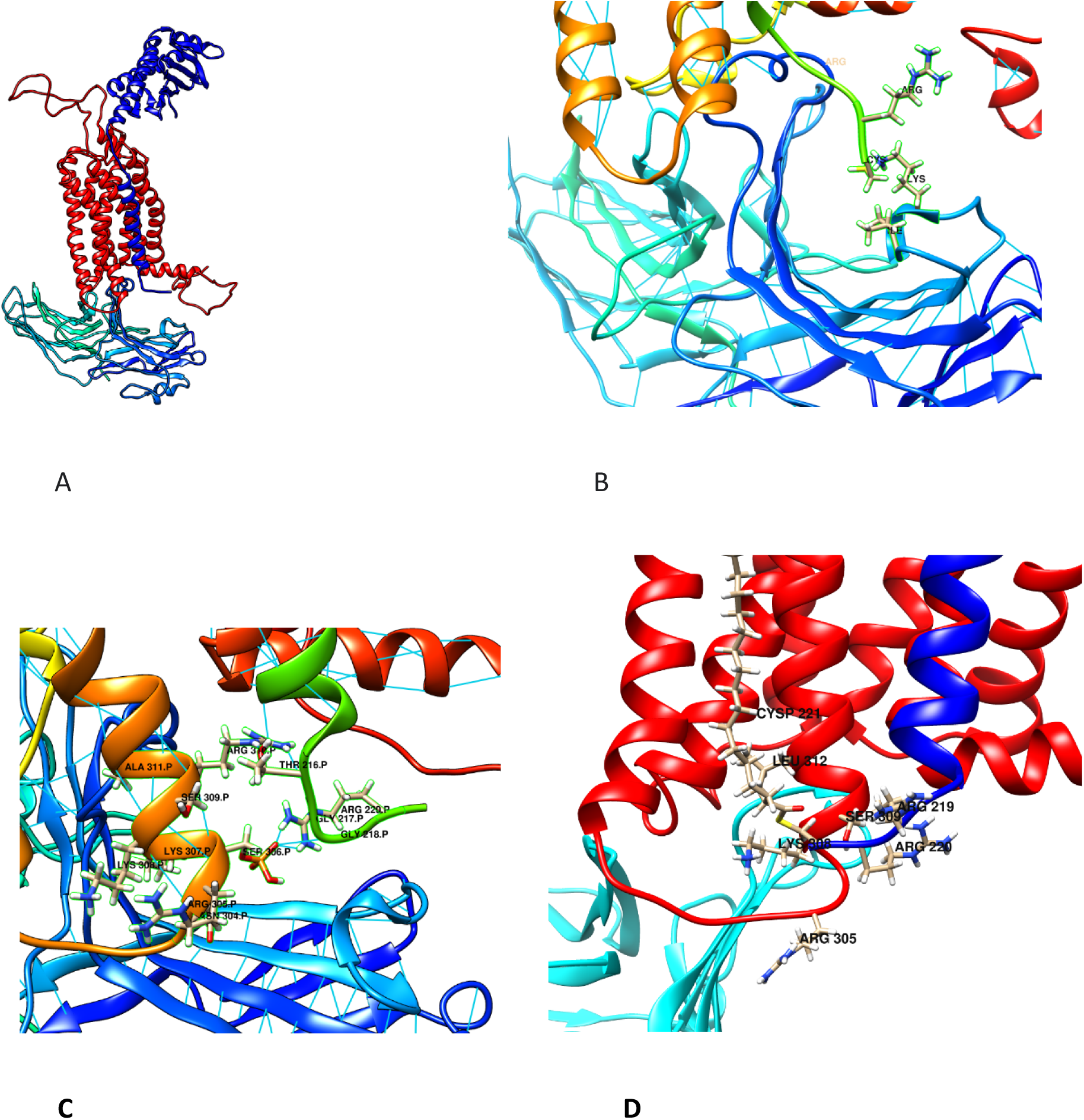
Interaction of the EPCR C-terminus with the PAR1-b-arrestin-2 (blue-cyan gradient) complex. **A**. The ternary PAR1 (red)-EPCR (blue) -β-arrestin-2 complex that initiates APC- induced PAR1 biased signaling. **B**. Possible direct interaction of depalmitoylated EPCR with b- arrestin-2. **C**. Interaction of depalmitoylated EPCR with b-arrestin-2 via involvement of the ICL3 and phosphorylated Ser306. **C**. Interaction of Cys221 palmitoylated (labeled as CYSP221 and colored by element) EPCR (blue-colored backbone with side chains colored by the element) with b-arrestin-2 (colored cyan) involving ICL3 and TM6 of PAR1 (red).

Changes in the FLR conformation are shown in figure 6B, while the rotation of N and C domains of b-arrestin-2 relative each other was evident from comparison of the initial and final time frames of the MD simulation (not shown). Our simulations were shorter (50 ns) than the simulations used in the literature for observing β-arrestin-2 activation, therefore changes we observed looked smaller than described in the literature. Figure 6B demonstrates a possible contact between Cys221 of EPCR and Lys residue in the N domain of b-arrestin-2. This observation raises the possibility that EPCR not only changes PAR1-β-arrestin-2 binding affinity by providing additional contacts, but also can help to activate the PAR1 bound β-arrestin-2 as seen from conformational changes in FLR. MD simulations of the PAR1-EPCR- β-arrestin-2 complex demonstrated that EPCR TMD facilitates PAR1 activation both via tethered peptide (figure 5C) and by the interaction of the 215-CTGGRRC-221 C-terminal sequence of EPCR with PAR1 (figure 6B, 6C, 6D). Because EPCR is palmitoylated and this process is reversible, we investigated both, palmitoylated and intact EPCR interaction with the PAR1-β-arrestin-2 complex. Docking of the 215-CTGGRRC-221 peptide to the β-arrestin-2 demonstrated binding of the peptide to the membrane facing region of β-arrestin-2. This raises the possibility that EPCR can directly interact with the β-arrestin-2 and add additional binding sites for β-arrestin- 2, thereby stabilizing the PAR1-EPCR-b-arrestin-2 complex (shown in figure 6B). Palmitoylated EPCR did not form direct contact with β-arrestin-2 but interacted with PAR1 residues involved in the interaction with the FLR (figure 6D). The overall effect of EPCR TMD on PAR1 includes the facilitated activation of PAR1 and stabilization of the PAR1-β-arrestin-2 complex.

## Conclusions

1. We reviewed published data on the initial stages of APC-biased PAR1 signaling and built three-dimensional structural models of individual proteins and protein complexes participating in the signaling. The models provide atomic-level interaction details consistent with the published biochemical observations and provide new insights into signaling events. The models allow us to propose plausible mechanistic explanations for some of the unexplained phenomena that make APC-biased PAR1 signaling distinctly different from thrombin-induced canonical PAR1 signaling.
2. Models of EPCR-APC interacting with the PAR1 N-terminus provide novel information on PAR1 residues involved in the interactions with the previously identified binding sites located in the catalytic domain and on the light chain of APC. In particular, the model explains that the preferred R46 cleavage by the APC-EPCR occurs when the 162-helix of APC binds to the PAR1 N- terminal peptide residues 22 to 32. This interaction places the APC catalytic domain into a position to favor the optimal cleavage of PAR1 R46 by the protease-receptor complex.
3. Models show that the length of even entirely extended conformation of PAR1 N-terminus will not allow it to interact with the binding sites in the catalytic domain and light chain at the same time. Although the models may not allow following all the steps in the time evolution of signaling events, models suggest the following sequence: first, cleavage of PAR1 peptide to form the TL, then interaction of TL with the APC light chain (possibly helping EPCR TMD to stay close to PAR1 TMDs), and then TL activation of the EPCR-PAR1 complex to initiate biased signaling.
4. The geometry of the PAR1 activation complex, including the length of TL and size of full- length models of EPCR and APC considered in the building of the PAR1 cleavage complex underline the importance of the hinge-like motions between extracellular and transmembrane domains of EPCR. These motions allow EPCR-bound APC to adjust active site position for effective cleavage of PAR1. The second insight derived from the geometry of the activation complex highlights the necessity of a certain distance between the TMD of EPCR and TMDs of PAR1 to enable cleavage of PAR1 by APC. This distance is approximately equal to the diameter of PAR1 or to the distance between binding sites for PAR1 and EPCR on the Cav1 oligomer. These distance relations may, at least in part, explain the transactivation of PAR1 by the EPCR- APC when PAR1 is in form of a homo- or heterodimer and also the Cav1 requirement for APC signaling.
5. The most influential assumption used in the model building was the consideration of direct binding between TMDs of PAR1 and EPCR and direct binding of both of these molecules to Cav1 oligomer. The possibility of direct binding follows from published co-isolation experimental data, while the structural and interaction details were derived from models. PAR1-Cav1 models suggest the existence of up to 11 binding sites per Cav-1 oligomer. EPCR-Cav1 models also suggest up to 12 binding sites per cav-1 oligomer. Taken together with the dependence of APC signaling but not thrombin signaling from Cav1, the models suggest that Cav1 can create a high local concentration of EPCR-APC and PAR1 positioned at the distance favoring the PAR1 activation by the EPCR-APC. This could, at least in part, explain previously unexplained different dependence of biased and canonical signaling from Cav1.
6. Comparison of Cav1-PAR1-EPCR-b-arrestin-2 models with the Cav1-PAR1-G protein models demonstrated Cav1 oligomer can block G protein-membrane binding in certain relative orientations of Cav1 and PAR1. Taken together with the ability of Cav1 to bind G proteins, the models suggest that Cav1 can provide a selective mechanism that favors b-arrestin-2-mediated biased signaling, and disfavors G proteins mediated canonical signaling.
7. Modeling assumptions about the direct binding of EPCR TMD to TM helices of PAR1 and investigation of binding details resulted in important observations which suggest that APC- biased signaling is actually EPCR-biased signaling. Published data suggest that biased signaling can be observed for PAR1 cleaved both at R41 or R46 by different proteases as long as proteases are bound to EPCR. This suggests that the ligand-biased PAR1 signaling paradigm should be more accurately described as co-receptor-biased signaling in the case of EPCR-APC signaling. Our models suggest that in this co-receptor-biased signaling, EPCR TMD bound to PAR1 to TM6, TM7, and intracellular helix 8 can facilitate PAR1 activation and help to stabilize b-arrestin-2 bound to activated PAR1.

## Methods or Models

### Molecular Dynamics simulations

The simulation systems for MD studies were created using the CHARMM-GUI web portal (19), with CHARMM36m additive force field, and with MD simulation parameters and input files tailored for NAMD2 software (20). In addition to the CHARMM36m force field for proteins and lipids (21), parameters for modified amino acids were generated using the CHARMM-GUI Input generator/Membrane builder (22). The modified amino acid residues participating in protein-protein or protein-lipid interactions included in the simulated systems were: γ-carboxyglutamic acid residues of APC Gla-domain, palmitoylated C- terminal Cys221 of EPCR, phosphorylated Ser306 of PAR1, EPCR extracellular domain-bound phosphatidylcholine ligand, and multiple disulfide bonds of APC and other molecules. The TIP3P explicit water model (23) was used for all simulations. The APC and β-arrestin-2 simulation systems contained water and ions to balance charges, while EPCR and PAR1 containing systems included membrane bilayer in addition to water and ions. The lipid content of the bilayer was similar to the caveolae and rafts and contained Cholesterol, Sphingomyelin, Phosphatidylcholine, Phosphatidylserine, and Phosphatidylethanolamine. MD simulation included minimization, equilibration in the NVT ensemble, and production runs in the NPT ensemble. Equilibration of the membrane bilayer-containing systems was carried out in six steps as suggested by the CHARMM-GUI Membrane builder software output. Production runs for all proteins and complexes were performed in ten 5 ns steps totaling 50 ns unless the duration of the simulation was changed as indicated in corresponding sections in the results.

Periodic boundary conditions (PBCs) were used in all simulations, and electrostatic interactions were calculated using the particle-mesh Ewald method (24) with a grid created automatically. Simulations were performed at constant pressure and temperature (303.15 K) (NPT ensemble) with a time step of 2 fs. Langevin dynamics was used along with algorithms implemented in NAMD2: Langevin piston method pressure control (25), and Rattle algorithm to enable a 2 fs time step (26). We performed the simulations using the Compute Unified Device Architecture (CUDA) version of NAMD2 or NAMD3 on the Lambda workstation with two Nvidia GeForce RTX 3090 graphical processing units (GPUs). The MD simulations in this work were used to refine structures built with molecular docking; to select the most likely conformations of the flexible protein segments not resolved in the published experimental structures; and most importantly, to reveal interaction details of the binding interfaces of newly modeled complexes.

### Protein structure prediction and homology modeling

Building full-length three-dimensional structures of the EPCR-APC, EPCR-Cav1, EPCR-PAR1, Cav1-PAR1, and PAR1-β-arrestin-2 complexes are described in the sections below. Briefly, the EPCR-APC complex was created by homology modeling using as a template experimental structure of the extracellular domain of the EPCR with bound Gla-domain of PC (PDB ID: 1lqv.pdb). The Cav1- EPCR and Cav1- PAR1 complexes were created by the molecular docking with the Hdock server (27) followed by the selection of the complexes that support known experimental observations. EPCR-PAR1 complex was created using the capability of the AlphaFold software to build homo- or hetero-oligomers. The AlphaFold software was used via a copy of the notebook AlphaFold2.ipynb which implements the protein folding algorithm. This notebook is available via Google Colaboratory (28). A model of the PAR1 N-terminus interacting with the APC was created using a combination of homology modeling and protein-peptide docking methods using the CABS-dock (29), Hpepdock (30), and HDOCK servers followed by the structure editing with UCSF Chimera software (31). The UCSF Chimera was also used for the analysis of structures and creation of images used in this manuscript. The VMD software (32) was used to analyze the MD trajectories and visualize Anisotropic Normal Mode (ANM) analysis results.

When templates for homology modeling were missing, the AlphaFold2 (28, 33) created highest scoring structures and complexes that meet the constraint requirements which were used for the study. The AlphaFold2 algorithm (34), uses multiple sequence alignment (35) to build three-dimensional structures from amino acid sequences and can create accurate complexes if there are thousands of similar sequences for alignment. In addition to experimental structures of proteins (Protein Data Bank; http://www.rcsb.org/pdb/) (36), this review uses the full-length protein models downloaded from the AlphaFold database (37–39).

### Anisotropic Normal Mode (ANM) analysis

The flexibility of proteins, relative motions of protein domains, and changes in domain orientation, not accessible at the time scales of the MD simulations, were studied using Anisotropic Normal Mode (ANM) analysis with ProDy software (40, 41).

## Acknowledgments

This work was in part funded by the NSF HBCU-UP RIA Catalyst Grant.

The author is grateful to Dr. Alireza Rezaie for reading this manuscript, and for his helpful critical comments.

